# Free-floating Bacteria Transcriptionally Respond to Shear Flow

**DOI:** 10.1101/2023.12.10.570978

**Authors:** Ashwin Ramachandran, Howard A. Stone, Zemer Gitai

## Abstract

Planktonic (free-floating) cells are typically assumed to be oblivious to any flow that carries them. Here we discover that planktonic bacteria can sense flow to induce gene expression changes that are beneficial in flow. Specifically, planktonic *P. aeruginosa* induce shear-rate-dependent genes that promote growth in low oxygen environments. Untangling this mechanism revealed that in flow, motile *P. aeruginosa* spatially redistribute, leading to cell density changes that activate quorum sensing, which in turn enhances oxygen uptake rate. In diffusion-limited environments, including those commonly encountered by bacteria, flow-induced cell density gradients also independently generate oxygen gradients that alter gene expression. Mutants deficient in this newly-discovered flow sensing mechanism exhibit decreased fitness in flow, suggesting that this dynamic coupling of biological and mechanical processes can be physiologically significant.

**One-Sentence Summary:** Free-floating bacteria integrate biological and mechanical responses to adaptively alter gene expression in flow environments

## Main text

Bacteria encounter complex and diverse mechanical environments in their lifestyle (*1*, *2*). For example, human pathogens can cause infections in environments with flow, including the bloodstream, urinary tract, and catheters (*3–5*). Previous studies have focused on the effects of flow that moves past surface-attached bacteria (*6–8*). However, the effect of flow on gene expression of bacteria that are being transported within the bulk of the flow has not been previously examined. Hydrodynamic considerations would suggest that in such environments, flow passively transports bacteria (*9–11*) and the shear stresses experienced by individual bacterial cells are smaller than those that the bacteria experience when swimming in the absence of flow (*12*). As a result, the general expectation is that planktonic bacteria are carried along with the flow as inert objects, and therefore, that shear flow would not alter the gene expression of planktonic (free-floating) bacteria (*13*). Here, we directly test whether bacteria respond to being transported by flow using the bacterium *Pseudomonas aeruginosa* (PAO1), a common pathogen implicated in cystic fibrosis, bacteremia, and urinary tract infections. Surprisingly, we find that flow does induce significant transcriptional responses in planktonic *P. aeruginosa.* Biophysical and molecular analyses reveal that this unexpected response emerges from a flow-dependent interaction between bacterial motility mechanics, cell signaling, and oxygen-related cellular processes under confined shear flow. This previously unappreciated bacterial response is likely to be physiologically significant, as we demonstrate that it promotes metabolic adaptation that is crucial for fitness in flow environments. Our findings also suggest that the strong coupling of physical and biological processes could help to understand other aspects of how bacteria adapt and thrive in physiologically important complex environments.

## Results

### Transcriptomics reveals flow-sensitive gene regulation in planktonic bacteria

To determine whether flow alters the gene expression of planktonic *P. aeruginosa,* we designed a long microfluidic channel that enabled us to expose bacteria to flow for significant lengths of time and then be fixed as they exit the channel. Since the shear rate varies across the channel cross-section, we characterized flow conditions in our experiments using the wall shear rate (see supplementary text). Using this microfluidic system, we performed bulk RNA-Seq analysis of planktonic bacteria that flowed for 55 min at a shear rate of 20 s^-1^, and compared gene expression of this population with a control population of planktonic bacteria under similar conditions with no flow (**Figure 1a**). We note that this shear rate is in the range that bacteria would encounter in environments like the bloodstream or urinary tract (*14*, *15*). Despite the fact that we expected that bacteria would not respond to the flow, our experiment revealed several genes that were differentially regulated between planktonic cells with and without flow (**Supplementary Table 1**). Among the most strongly flow-induced genes were several gene clusters that were previously associated with the response to low oxygen conditions (*16–18*), including *nar*, *arc*, and *hcn* (**Figure 1b**, **Supplementary Table 1**).

**Figure 1.**
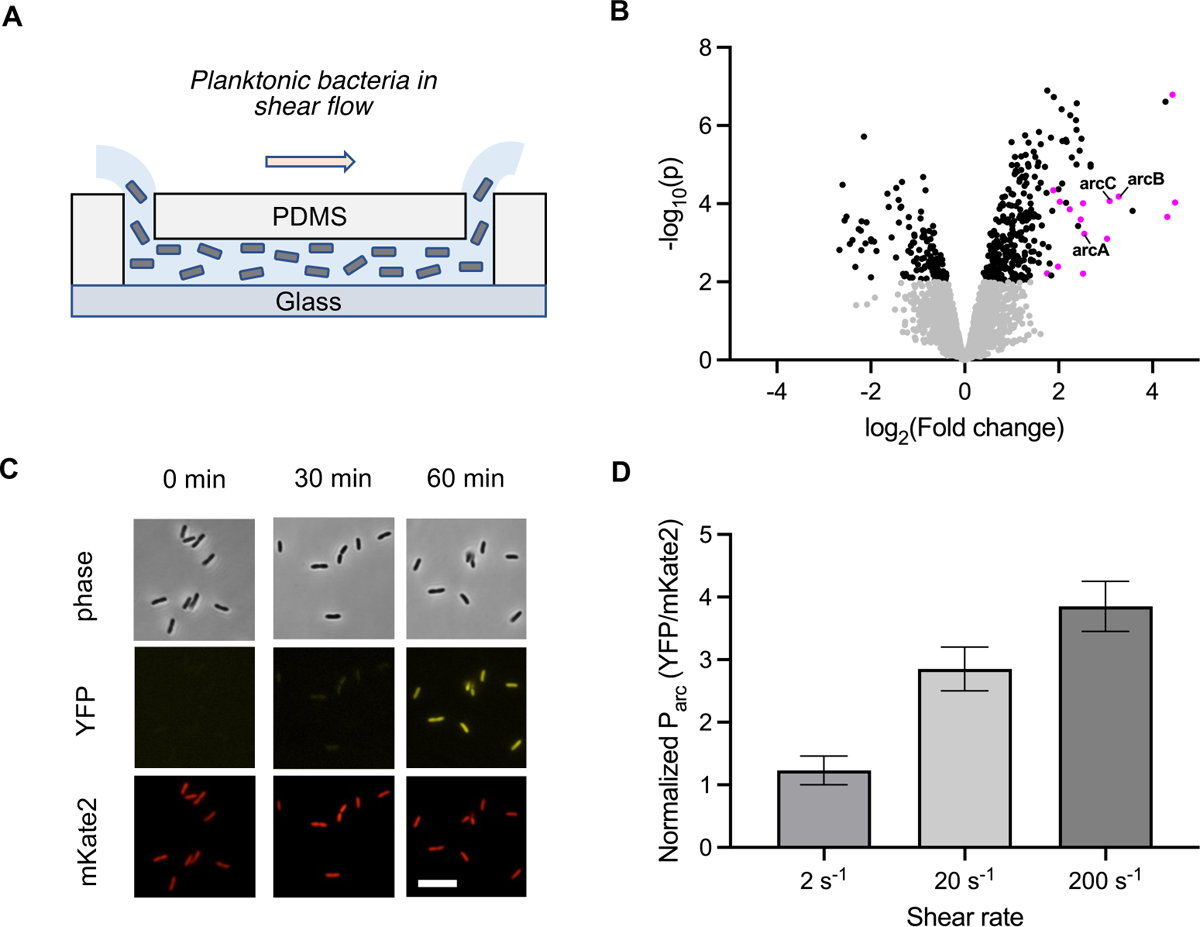
Flow induces transcription of low-oxygen genes in planktonic *P. aeruginosa*. (**A**) Schematic of microfluidic system used in this work to study the interaction between planktonic bacteria and shear flow. (**B**) Volcano plot representing significance versus fold-change of transcripts from bulk RNA-Seq of *P. aeruginosa* that flowed in the channel for 55 min at 20 s^-1^ compared to transcripts from cells under no flow conditions within the channel for the same duration. A *p*-value of 0.01 is used as the threshold for significance. Symbols in magenta represent genes that are upregulated under low oxygen conditions (see **Supplementary Table 1** for a complete list of these genes and for all the genes that are upregulated by at least three-fold due to flow). Also indicated by solid lines are genes from the *arc* operon. (**C**) Representative phase contrast and fluorescence microscopy images of cells after subjecting them to shear flow of 20 s^−1^ for 0, 30, and 60 min. YFP and mKate2 represent transcriptional reporter signals from individual cells corresponding to the transcription of the *arc* operon and constitutively expressed *rpoD*, respectively. Scale bar: 8 μm. (**D**) Relative fold change of *arc* transcriptional signal for flow at 2, 20, and 200 s^-1^ compared to no-flow conditions. The transcriptional signal for each condition was obtained as the normalized ratio of YFP to mKate2 fluorescence from individual cells, averaged across at least 30 cells from three independent experiments. Error bars represent the s.d. from the three replicates and each flow condition is significantly different from the other with a threshold *p* value 0.01. Statistical significance is calculated using unpaired *t*-tests.

To study flow-dependent transcriptional responses in single cells and to monitor the dynamics, we engineered an *arc* transcriptional reporter PAO1 strain in which the *arc* operon promoter is fused to YFP and a constitutive *rpoD* promoter (*19*) is fused to mKate2 for normalization. Consistent with our RNA-Seq data, we observed that the *arc* reporter signal significantly increased with time under flow for over 1 h (**Figure 1c**). To determine the quantitative relationship between flow rate and *arc* induction, we examined the *arc* reporter under flow after 1 h at shear rates of 0 (no flow), 2, 20, and 200 s^-1^. We observed that the induction of the *arc* reporter increased monotonically with shear rate, suggesting that planktonic bacteria under shear flow exhibit a low-oxygen transcriptional response in a flow-sensitive manner (**Figure 1d** and **Supplementary Figure 1**).

### Shear flow causes swimming bacteria to redistribute and form cell density gradients

How might flow affect planktonic bacteria? To gain insight into the mechanism underlying the flow-dependent *arc* induction we examined the physical response of planktonic bacteria to shear flow in a defined rich media, EZ. Specifically, we performed phase contrast microscopy to image planktonic bacteria in shear flow in the middle of the channel (away from the surface). We observed that these bacteria distributed to create spatial gradients with more cells concentrated towards the channel walls and fewer cells concentrated towards the channel center (**Figure 2a**). We quantified this response at varying shear rates and found that the cell depletion near the channel center and the corresponding cell enrichment near the walls both increased with shear rate (**Figure 2b** and **Supplementary Figure 2**). For example, at a shear rate of 20 s^-1^, the cell density decreased by ∼2-fold near the channel center and increased by ∼2-fold near the channel walls compared to no-flow conditions.

**Figure 2.**
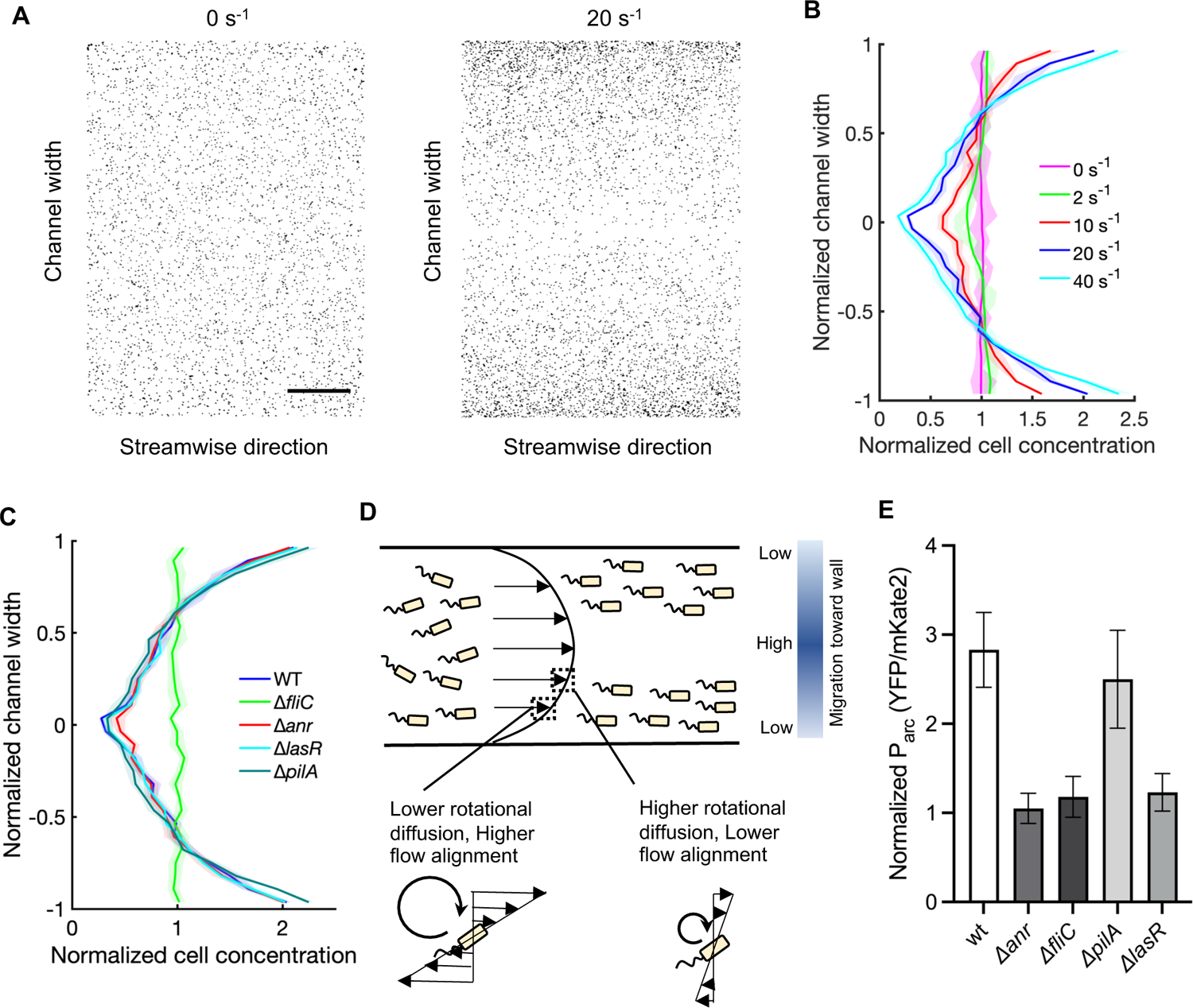
Flagellar motility causes planktonic bacteria to form cell density gradients in flow, and flow-mediated *arc* induction requires flagellar motility, quorum sensing, and oxygen sensing. (**A**) Instantaneous cell distribution of WT *P. aeruginosa* measured across the channel width under no-flow (0 s^-1^) and flow at a shear rate of 20 s^-1^. Five images were taken for each flow condition. Scale bar: 250 μm. (**B**) Cell density distribution across the channel width averaged within the field of view at shear rates of 0, 2, 10, 20, and 40 s^-1^. The solid line and shaded regions for each condition represents the mean and standard deviation of the distribution from across five independent measurements. The channel width is normalized by the half-width of the channel. (**C**) Cell density distribution across the channel width averaged within the field of view for WT, Δ*anr*, Δ*fliC*, Δ*pilA*, and Δ*lasR* strains for flow at a shear rate of 20 s^-1^. (**D**) Schematic of the shear-trapping model. (**E**) Normalized *arc* transcriptional reporter signal for WT, Δ*anr*, Δ*fliC*, Δ*pilA*, and Δ*lasR* strains for flow at a shear rate of 20 s^-1^, averaged across at least 30 cells from three independent experiments. Error bars represent the standard deviation.

Previous studies have suggested that hydrodynamic interactions between self-propulsion motility and shear gradients could affect the distribution of bacteria in flow (*13*, *20*). We thus tested the effect of Type IV pilus (T4P) and flagellum-dependent motility mutants on our flow-dependent redistribution pattern. We found that flagellar mutants no longer established cell density gradients, while the behavior of pilus mutants resembled that of wild type PAO1, suggesting that flagellar motility is required for shear-induced cell redistribution (**Figure 2c** and **Supplementary Figure 3**). Our observations on cell redistribution in flow are consistent with previous studies on the effects of flow on rod-shaped bacteria with active swimming motility (*21*). This transverse redistribution of swimming bacteria in response to shear gradients has been modeled as a “shear-trapping” mechanism (**Figure 2d**) (*21–23*). In the shear-trapping model, bacteria near the channel walls encounter a transverse diffusive-like redistribution from the velocity gradient that aligns the cell body along the flow direction, and this counteracts rotational noise from swimming. This effect causes bacteria that are closer to the walls to remain more aligned with the flow and thus remain close to the wall, while bacteria closer to the channel center (low shear rates) are more likely to swim away from the center. In this manner, flow causes bacteria to tend to migrate away from the center of the channel and get trapped near the walls over a short diffusive timescale (*22*, *23*).

A key feature of the shear-trapping model is that it depends on the velocity gradient, which is influenced by the shear rate but not the shear stress. To test if the shear-trapping model is likely to explain the cell density gradients we observed, we varied the viscosity of the cell media because increased viscosity increases the shear stress without increasing shear rate for a given flow rate. We found that the bacterial cell density distribution in flow did not change significantly upon increasing viscosity (**Supplementary Figure 4**), confirming that the velocity gradient is the primary driver in setting the cell density profiles in flow.

### The transcriptional flow response requires flagellar motility, quorum sensing, and oxygen sensing

The experiments above demonstrate that flow induces both transcriptional induction of *arc* and cell density changes. Furthermore, the density changes require flagellar motility and *arc* is known to be regulated by the Anr oxygen sensing system (*24*). To understand how each of these processes affect *arc* induction in flow, we assayed flow-dependent *arc* induction and cell density redistribution in mutants that disrupt cell density sensing (the quorum sensing master regulator, *lasR* (*25*, *26*)), cell motility (*fliC* for flagellar motility and *pilA* for T4P (*27*, *28*)), and oxygen sensing (*anr*) (*24*, *29*)*. pilA* had no effect on cell density or *arc* induction compared to wild-type (WT), both *anr* and *lasR* deletion cells resembled WT with respect to cell density gradients but failed to induce *arc* in flow, and *fliC* was defective in both cell density redistribution and *arc* induction (**Figure 2c**, **2e**, and **Supplementary Figure 3**). Based on these data we hypothesized that flagellar motility is required for establishing cell density gradients in response to flow and that the resulting cell density gradients subsequently induce *arc* expression through the LasR and Anr regulators.

### Bacterial redistribution in flow leads to increased oxygen uptake rate via cell density sensing

The dependence of *arc* induction in flow on motility-driven cell gradient formation suggested that the cell density gradient that occurs in flow might produce an oxygen concentration gradient that activates Anr. To test this hypothesis, we obtained *in situ* measurements of oxygen concentration in the middle of the channel by measuring the fluorescence of an oxygen-sensitive dye dissolved in EZ media after 1 h of shear flow (see supplementary text). Counter to our hypothesis, we did not observe an oxygen gradient in these conditions, suggesting that there is an alternative mechanism explaining *arc* induction in flow (**Figure 3a**). For example, if flow caused bacteria to consume oxygen at a higher rate, the overall level of oxygen would decrease more quickly in flow than in the absence of flow, causing Anr to be activated by flow despite the absence of a spatial oxygen gradient. Consistent with this idea, our oxygen sensor revealed that overall oxygen levels decreased in flow faster than in the absence of flow over the course of one hour in EZ media (**Figure 3b**). We then quantified oxygen uptake rate by averaging the measured oxygen concentration across the field of view and comparing this average value measured at different locations along the channel (equivalently, in time). Under flow, the wild-type PAO1 consumed oxygen at a rate that was ∼30% higher than the no-flow condition (**Figure 3c**).

**Figure 3.**
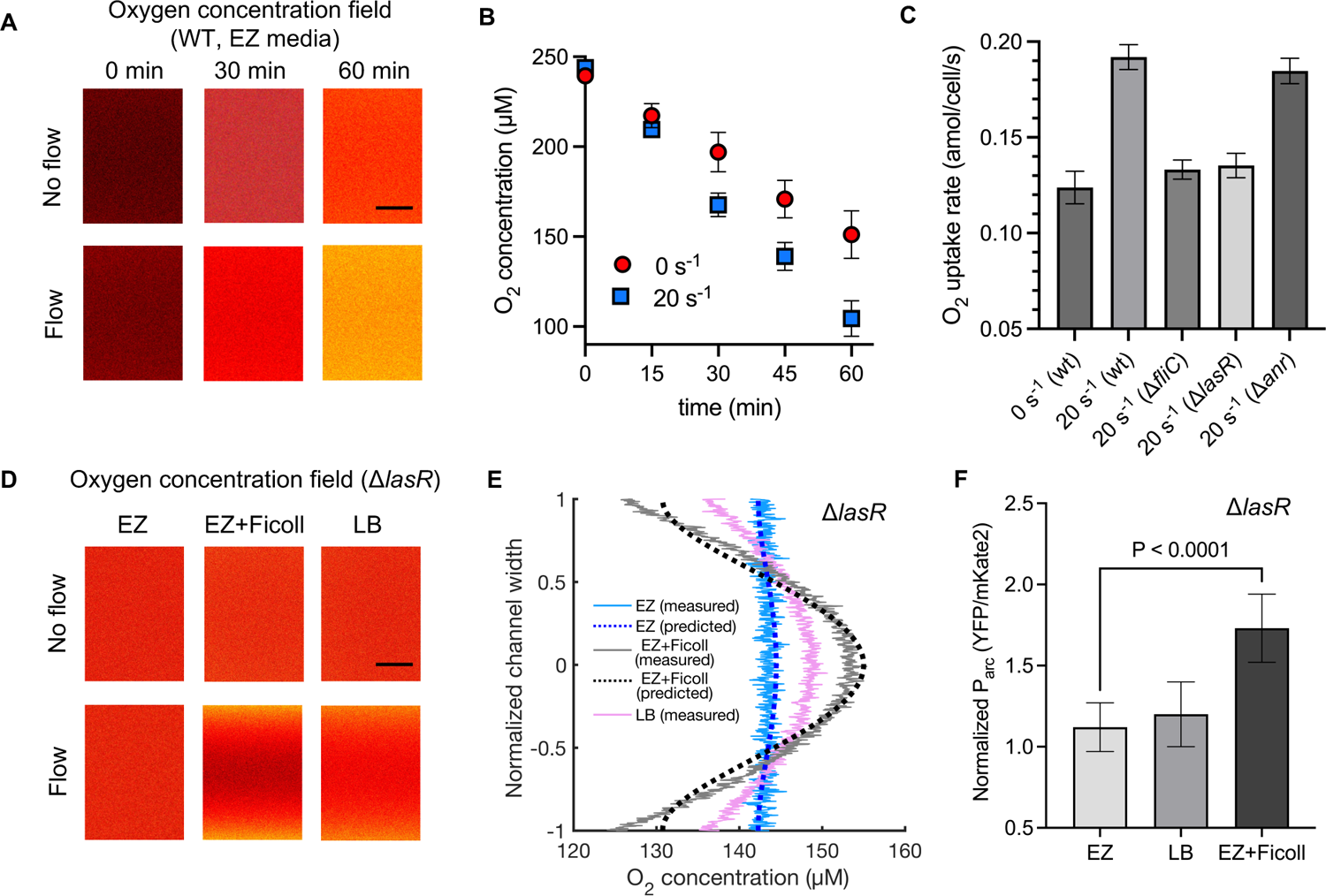
Quorum sensing increases oxygen uptake rate in flow, and spatial oxygen gradients form under diffusion-limited conditions. (**A**) Heatmaps depicting measured oxygen concentration fields within the channel versus time for no flow and flow at 20 s^-1^ of planktonic WT *P. aeruginosa* in EZ rich media, measured using RTDP fluorescent dye dissolved in the media (see supplementary text). RTDP fluorescence is quenched by oxygen, so an increase in fluorescence indicates reduced oxygen concentration. Scale bar: 400 μm. (**B**) Average oxygen concentration within the field of view versus time for WT *P. aeruginosa* in EZ rich media for no flow and flow at 20 s^-1^. Plotted are the mean values from three independent experiments and the error bars represent the standard deviation. (**C**) Average oxygen consumption rate for no flow conditions of WT and for flow at a shear rate of 20 s^-1^ of planktonic WT, Δ*anr*, Δ*fliC*, and Δ*lasR* strains. Shown are the mean values and standard deviation from three independent experiments. (**D**) Heatmaps of measured oxygen concentration fields within the channel after 1 h of no flow and flow at 20 s^-1^ of Δ*lasR* strain in EZ, EZ + 15% Ficoll, and LB media. Scale bar: 400 μm. (**E**) Measured oxygen concentration (solid lines) averaged across the streamwise direction within the field of view versus channel width after 1 h of flow at shear rate of 20 s^-1^ of Δ*lasR* strain in EZ, EZ + 15% Ficoll, and LB media. Dashed lines represent predictions based on the reaction-diffusion model. The channel width is normalized by the half-width of the channel. (**F**) Normalized *arc* transcriptional reporter signal for Δ*lasR* strains for the same conditions as (**E**), averaged across at least 30 cells from three independent experiments. Error bars represent the standard deviation.

Why do *P. aeruginosa* cells consume oxygen at higher rates under flow? We wondered if this increased uptake rate of oxygen in flow was related to the other transcriptional regulator we found to affect flow-dependent *arc* induction, the LasR quorum sensing master regulator. Indeed, the oxygen consumption rate of Δ*lasR* mutants was not flow dependent and resembled the oxygen consumption rate that WT exhibited under no flow conditions even when Δ*lasR* was exposed to 20 s^-1^ flow for one hour (**Figure 3c** and **Supplementary Figure 5**). Similarly, loss of flagellar motility, which eliminated the cell density redistribution, also caused the bacteria to lose the flow dependence of oxygen consumption. Meanwhile, *anr* mutants maintained flow-dependent oxygen consumption similar to that of WT (**Figure 3c** and **Supplementary Figure 5**). These results suggest that motility-dependent changes in bacterial cell density activate quorum sensing through LasR and that LasR induction increases the oxygen consumption rate.

LasR has been previously implicated in a number of metabolic changes that could alter oxygen consumption rate in a cell-density-dependent manner (*16*, *18*, *30–32*). To test if the regulation of oxygen uptake rate by *lasR* can be driven by quorum sensing independently of flow, we quantified the oxygen uptake rate in WT and Δ*lasR* strains using spent media or increased concentration of cells within the same media under no flow conditions. We observed that only the wild-type strain showed an increase in oxygen uptake rate in both conditions, suggesting that the regulation of oxygen uptake rate via *lasR* is a flow-independent response (**Supplementary Figure 6**). Together, these data suggest that shear flow of planktonic bacteria leads to an overall increase in oxygen uptake rate, and this requires the coupling of swimming motility with the ability to sense cell density changes in flow.

### Diffusion-limited conditions can lead to oxygen gradients in flow

To better understand oxygen sensing at a quantitative level we developed a reaction-diffusion model to describe how flow affects oxygen concentration fields in the presence of cell gradients. This model suggested that spatial oxygen gradients should emerge when oxygen diffusion is slower than the oxygen consumption rate (**Supplementary Figure 7**). To test if the absence of an oxygen gradient observed in WT PAO1 in EZ media is due to the rapid diffusion of oxygen in these conditions, we decreased oxygen diffusion 10-fold by increasing the viscosity of the EZ media with added Ficoll (*7*) (the diffusion versus viscosity correspondence follows the Stokes-Einstein relation). To focus on changes in oxygen diffusion rather than effects on oxygen consumption rate, we performed these experiments in Δ*lasR* mutants. As predicted by the model, we found that Ficoll supplementation caused these bacteria to produce significant oxygen gradients, as measured by our oxygen sensor dye (**Figures 3d** and **3e**). Moreover, we also discovered that in these diffusion-limited conditions, *arc* induction occurred even in the absence of *lasR* (**Figure 3f**). These results suggest that motility-dependent cell density changes in flow can sustain spatial oxygen gradients that are sufficient to alter bacterial gene expression when oxygen diffusion is limited.

We were surprised to discover that unlike in EZ media, oxygen gradients were sustained in LB media even in the absence of Ficoll (**Figures 3d** and **3e**). To quantify the effect of LB on oxygen diffusion we turned to our validated reaction-diffusion model. The measured oxygen diffusion coefficient in water at room temperature is 2×10^-9^ m^2^/s. Using this value accurately captures our experimentally-measured oxygen gradient profile in EZ media without Ficoll, and reducing this value 10-fold accurately captured the gradients measured in EZ media with Ficoll supplementation (**Figure 3e**). However, this oxygen diffusion coefficient failed to fit our data for LB. Varying the oxygen diffusion coefficient value revealed that the model fit was best with an oxygen diffusion coefficient of 0.7×10^-9^ m^2^/s (**Supplementary Figure 8**). The viscosity of LB and EZ media are similar, suggesting that their difference in oxygen diffusion could result from oxygen’s interactions with the complex macromolecules found in LB but absent from EZ. We tested this by adding tryptone and yeast extract (complex components of LB) individually to EZ media in concentrations typically used for LB. In both cases, we observed that adding these components produced small gradients in oxygen concentration across the channel width, suggesting that macromolecular components in LB media tend to slow the diffusion of oxygen (**Supplementary Figure 8**).

### Oxygen sensing promotes planktonic bacterial fitness in flow

Our observations that planktonic bacteria induce a transcriptional response to flow raise the question of whether this transcriptional response provides a fitness advantage. To assess the potential fitness benefits of flow-induced transcription we compared the growth rates of planktonic WT PAO1 and mutants defective in *arc* induction in flow. Specifically, we monitored the growth of each strain after subjecting the cells to flow at varying shear rates for over 2.5 hours (roughly two doubling times). In WT PAO1 we observed no growth rate changes as a function of flow. We next assayed growth in Δ*anr* mutants, as these cells redistribute to form cell gradients and exhibit flow-induced changes in oxygen uptake, but are defective in oxygen sensing. We observed that the Δ*anr* strain showed a significant decrease in growth rate at higher shear rates (**Figure 4a**), suggesting that there is an adaptive benefit to oxygen sensing for planktonic cells in flow. In contrast, Δ*fliC* cells, which do not redistribute and thus do not preferentially deplete oxygen in flow, and Δ*lasR* cells, which do not increase oxygen uptake and thus also do not preferentially deplete oxygen in flow, had similar growth rates across all flow rates. These data suggest that WT PAO1 utilizes the Anr oxygen sensing pathway to help it adapt to the low oxygen environment induced by flow’s effects on motile planktonic cells.

**Figure 4.**
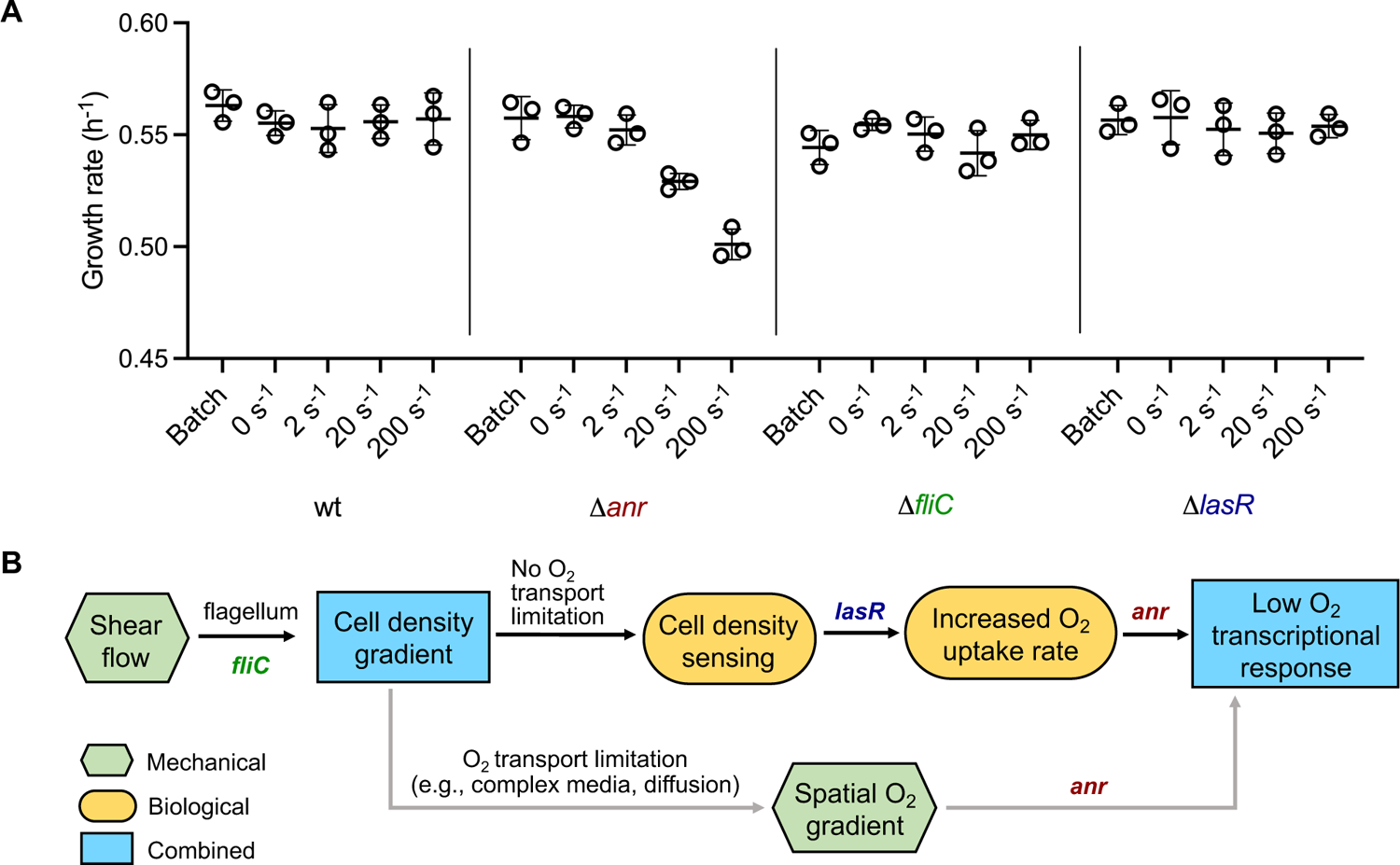
Oxygen sensing provides a fitness advantage to planktonic bacteria in flow. (**A**) Growth rate measurements of planktonic WT, Δ*anr*, Δ*fliC*, and Δ*lasR* strains of *P. aeruginosa* strains in batch culture and at varying shear rates of 0, 2, 20, 200 s^-1^. Shown are individual values from three independent experiments and indicated are the mean and standard deviation. (**B**) Biophysical mechanistic model for the interaction between shear flow and planktonic bacteria. Shown in green hexagons are factors that are derived purely from mechanics. A purely biological mechanism is indicated by the yellow ellipses. Blue rectangles indicate phenomena that result from combined mechanical and biological mechanisms. At each step in the mechanistic model, the corresponding genes or pathways are indicated. The black arrows represent a mechanism where there is an increase in oxygen uptake rate resulting from the coupling of flagellar motility, quorum sensing, and oxygen sensing. The gray arrows indicate a mechanism that occurs when oxygen diffusion is limited, independent of quorum sensing.

## Discussion

Here we present data suggesting a biophysical model for how interactions between hydrodynamics and bacterial processes like motility, metabolism, and signaling cause planktonic bacteria to alter their gene expression to provide adaptive benefits in the presence of shear flow (**Figure 4b**). First, shear flow causes bacterial cells to redistribute and become more crowded near the channel walls via shear trapping, which results from the interaction of flagellar motility and hydrodynamics of shear flow. Under conditions where oxygen diffusion is not limited, cells sense their density changes in flow via quorum sensing, causing them to increase their oxygen uptake rate. After time, the resulting increased oxygen uptake reduces the overall oxygen concentration in the flow environment. This lower oxygen concentration is sensed by Anr, resulting in flow-induced upregulation of genes associated with microaerobic and anaerobic conditions. Independent of quorum sensing, if the cells are present in an environment where oxygen diffusion is limited, the greater oxygen consumption resulting from the higher cell density near the walls is sufficient to establish a spatial oxygen gradient, resulting in low-oxygen niches and an Anr-mediated transcriptional response. This newly-discovered ability of planktonic bacteria to adapt their gene expression profiles in flow may be functionally significant because mutants that cannot perform this adaptation exhibit compromised fitness in flow.

Our study highlights how the coupling of bacterial mechanics in shear flow and regulation of cell density-dependent metabolic processes can lead to unexpected behaviors of bacterial populations in complex environments. We note that a purely mechanics-based consideration would have predicted that planktonic bacteria should simply move with the flow and thus not have any signals available to sense the presence of flow. The finding that the bacterial motility and signaling pathways enable them to sense and respond to flow thus illustrate the complex behaviors that can emerge when living systems interact with their mechanical environments.

While the conditions we tested here used a simple channel geometry and standard culture media, our study also points to the possibility of diverse interactions between populations of planktonic cells and flow in physiologically relevant contexts. For example, most current studies assume that the diffusion of oxygen in biological systems is similar to the diffusion of oxygen in water. However, our observation that oxygen diffusion is limited in media with complex solutes suggests that biological fluids such as blood and plasma can present constraining environments for the free diffusion of oxygen (*33*). This finding thus calls for a careful reevaluation of oxygen diffusion in biological systems instead of assuming a standard value across all media compositions and conditions.

While our study was primarily focused on oxygen transport and metabolism under flow, the motility-dependent cell density gradients established in flow could also lead to concentration gradients of other molecules such as nutrients, signaling molecules, proteins, and secondary metabolites. Indeed, these molecules all diffuse more slowly than molecular oxygen, so they would be predicted to form even stronger gradients (**Supplementary Figure 7**). This further points to the possibility that cells are likely to encounter a complex spatially stratified biochemical landscape under confined shear flow, which would not be seen in standard laboratory growth conditions. The effect of such a complex biochemical landscape around bacteria in shear flow on physiological fitness and adaptation over both short and long-timescales remains to be explored in detail.

Given how intricately the swimming motility of bacteria is coupled with the biophysical and molecular response under flow, it remains to be seen how other cell types respond when they are carried along by flow. For example, cells with distinct shapes or motility mechanisms will have different hydrodynamic interactions with shear flow that could result in different emergent behaviors. Beyond examining how additional bacterial and eukaryotic species respond to flow individually, it will prove interesting to understand how flow affects the biophysical and biochemical interactions of planktonic polymicrobial communities like those of free-floating microbiome bacteria in the lumen and mucosa of the human gastrointestinal tract (*34*). Thus, our findings that bacteria are not merely transported by flow but rather actively respond to this complex environment presents an important step towards understanding the interconnected ways by which the coupling of biology and mechanics can produce surprising yet physiologically-significant adaptations.

## Acknowledgements

We thank Joshua W. Shaevitz, members of the Gitai Lab, and members of the Shaevitz lab for helpful discussions and feedback. **Funding:** This work was supported by the National Science Foundation grant MCB 2033020 (to ZG and HAS). **Author contributions:** All authors conceptualized and developed the methodologies of the study, AR performed experiments and data analysis, all authors contributed to data interpretation and writing the manuscript. **Competing interests:** Authors declare that they have no competing interests. **Data and materials availability:** All of the RNA-Seq data used to reach the conclusions of this paper are freely available under the National Center for Biotechnology Information Sequence Read Archive submission number <u>PRJNA1049796</u>. All other data are available in the main text or the supplementary materials. MATLAB codes for the numerical model and image analysis are available from the corresponding author upon request.

## Supplementary Materials for

### Materials and Methods

#### Bacterial strains and culture conditions

A list of the strains used in this study can be found in **Supplementary Table 2**, the plasmids used are described in **Supplementary Table 3**, and the primers used are described in **Supplementary Table 4**.

*P. aeruginosa* PAO1 was grown overnight in liquid EZ rich defined medium (*35*) (Teknova) or Luria-Bertani media (LB) Miller (Difco) in a floor shaker at room temperature of 25°C. Prior to flow experiments, overnight-grown cultures were diluted 1:200 in fresh media and grown to mid-exponential phase (OD of 0.4-0.5). For experiments involving Ficoll, cells were first grown to mid-exponential phase as described above in media without Ficoll, and when the cells reached the desired OD, the media was supplemented with 15% Ficoll prior to flow.

For cloning, PAO1 was grown on LB Miller agar (1.5% Bacto Agar) and on Vogel-Bonner minimal medium (VBMM) agar (200 mg/l MgSO4.7H2O, 10 g/l K2HPO4, 2 g/l citric acid, 3.5 g/l NaNH4HPO4.4H2O, and 1.5% agar) at 37°C, and on no-salt LB (NSLB) agar (5 g/l yeast extract, 10 g/l tryptone, 5% sucrose, and 1.5% agar) at 30°C. *Escherichia coli* S17 was grown in liquid LB Miller (Difco) in a floor shaker and on LB Miller agar (1.5% Bacto Agar) at 30°C or at 37°C. Antibiotics were used at the following concentrations: 200 μg/mL carbenicillin in liquid (300 μg/mL on plates) or 10 μg/mL gentamycin in liquid (30 μg/mL on plates) for *Pseudomonas*, and 100 μg/mL carbenicillin in liquid (100 μg/mL on plates) or 30 μg/mL gentamycin in liquid (30 μg/mL on plates) for *E. coli*.

#### Strains and plasmid construction

PAO1 deletion strains were constructed following the two-step allelic exchange protocol (*36*) using the plasmid pEXG2. Fragments ∼500 bp directly upstream and downstream of the target gene were amplified from WT PAO1 using primer pairs P1/P2 and P3/P4 (**Supplementary Table 4**), respectively. The resulting upstream and downstream fragments were fused using overlap-extension PCR with primer pair P1/P4. The resulting fragment was cloned into the HindIII site of plasmid pEXG2. The pEXG2 plasmid was then integrated into *P. aeruginosa* PAO1 through conjugation with the donor strain *E. coli* S17. Mating was performed on LB plates, and the exconjugants were selected on VBMM plates containing 30 μg/mL gentamycin. Mutants of interest were counter-selected on NSLB plates supplemented with 15% (w/v) sucrose. Several single colonies were screened for the correct mutation using PCR and amplicon sequencing (SNPsaurus) with the primer pair P1/P4.

For constructing *arc* transcriptional reporter strains of PAO1, the fragment containing the *PaQa* promoter region within the plasmid pPaQa (*19*) was replaced with a fragment containing the *arc* operon promoter. First, the plasmid pPaQa was digested and linearized using restriction enzymes XhoI and BamHI to remove the *PaQa* promoter region. Thereafter, a region ∼490 bp upstream of the start codon of the *arc* operon which contained the *arc* promoter was PCR amplified, and the resulting amplicon together with the digested plasmid were Gibson assembled. The resulting plasmid was electroporated into *E. coli* S17 for maintenance. Selective media containing carbenicillin was used for the plasmid maintenance. Plasmids were then purified from the S17 maintenance strain using the Qiagen miniprep kit and then introduced into PAO1 using electroporation.

#### Microfluidic system design and flow conditions

Microfluidic channels were custom-fabricated using polydimethylsiloxane (PDMS) bonded to large glass slides (Ted Pella, USA). Molds for PDMS channels were 3D printed on an ABS-like WaterShed XC 11122 material (Proto Labs, USA). The channel had a rectangular cross-section with a depth and width of 425 μm and 1500 μm, respectively. The channel length followed a serpentine geometry with 34 straight sections that were each 150 mm long, corresponding to a total channel length of approximately 5 m. Flow conditions were characterized using the wall shear rate, given by:

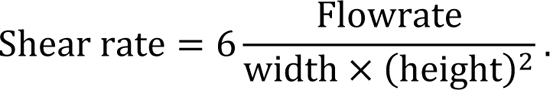

Our channel geometry enabled bacteria to be transported in flow within the channel for up to 1 h at a shear rate of 20 s^-1^. A 6 mL bacterial suspension in media was loaded completely into a 10 mL syringe and the solution was injected into the channel using a syringe pump. The flow rate was adjusted according to the desired shear rates. For experiments at shear rate of 200 s^-1^, a peristaltic pump was used in place of syringe pump to recirculate bacteria within the channel and enable long-duration exposure of bacterial cells to flow. All flow experiments were performed at room temperature.

#### RNA-Seq of planktonic bacteria in shear flow

For the RNA-Seq experiments, a branched channel section near the device outlet was used to inject 37% Formaldehyde at a flow rate equal to 1/8th of the main channel to fix planktonic bacteria in a 4% Formaldehyde (final concentration) media solution after they flowed for 55 min within the device and prior to exiting the channel. This allowed us to collect a sufficient number of fixed bacterial cells required to perform transcriptomics on a population that was exposed to flow for a set duration. For the no-flow control, bacterial cells were incubated for 55 min under static flow conditions within the channel and then pipetted out and fixed for the same duration as flow-exposed cells.

RNA was extracted from fixed cells using the Qiagen RNeasy Mini kit with the following modifications to the protocol. Fixed cells were spun down at 6000 ×g for 5 min and resuspended in an 80 µL volume containing 30 mM Tris-HCL (pH 8.0), 2 mM EDTA, and 0.1% Triton-X 100, and 20 mg/mL lysozyme. Lysis was performed at 37°C for 20 min, followed by Proteinase K digestion at 65°C for 1 h. In addition, on-column DNase digestion was performed during RNA extraction as per the Qiagen RNeasy protocol. The extracted total RNA was then sent to SeqCenter (Pennsylvania) who generated libraries using Illumina Stranded RNA library preparation and RiboZero Plus rRNA depletion. Sequencing was performed by SeqCenter on an Illumina NextSeq2000 platform providing up to 12 million paired end reads (2×51 bp) per sample. After sequencing, SeqCenter conducted an intermediate RNA-analysis and provided a list of differentially expressed genes. Briefly, quality control and adapter trimming were performed with bcl2fastq, read mapping was performed with HISAT2, and read quantification was performed using Subread’s featureCounts. Thereafter, read counts were normalized using edgeR’s Trimmed Mean of M values algorithm and subsequently converted to counts per million. Differential expression analysis was performed using edgeR’s Quasi-Linear F-Test (qlfTest) functionality against treatment groups, with threshold of |log2(fold-change)| > 1 and *p*-value < 0.01.

#### Cell density measurements in flow

Phase contrast microscopy was used to measure the cell distribution of planktonic bacteria in shear flow. Imaging was performed using a 10x objective on a Nikon Eclipse Ti microscope connected to a digital CMOS camera (Hamamatsu ORCA-FLASH4.0). A short exposure time corresponding to the flowrate was used to avoid cell streaking and minimize scatter from the channel edges. The imaging region was chosen to be the middle of the channel cross-section, i.e., halfway deep, away from the walls at a location approximately 4 m from the channel inlet and the region of interest spanned the entire channel width. Between successive measurements at different flowrates, at least 10 min was provided for the cell density distribution to equilibrate. Thereafter, five images were taken at a given shear rate in 1 min intervals. The cell density distribution was obtained by averaging across these measurements. A custom MATLAB script was developed to calculate the cell density distribution from experimental images. Briefly, the raw images were denoised, background subtracted using a threshold, and binarized to identify individual bacterial cells and their corresponding centroids. Thereafter, the channel width was binned into 100 sections, and within each section the total number of cells was computed and normalized by the total number of cells within the entire field of view. The resulting normalized value of cell counts was plotted versus the bin centers to obtain the cell density distribution.

#### Transcriptional reporter assays

For the transcriptional reporter assays, the reporter strains were first subject to flow at varying shear rates and durations within the channel. The high speeds of bacteria during flow within the channel precluded direct fluorescence imaging of the cells during flow. Therefore, after subjecting the cells to a set duration in flow, the cell solution from the channel was immediately collected in a 1.5 mL Eppendorf tube and a ∼10 μL volume of the solution was sandwiched between a cover glass and glass slide and imaged. Phase contrast and fluorescence (YFP, mKate2) microscopy was performed on a 100x/1.4 NA objective on a Nikon Eclipse Ti microscope connected to a Hamamatsu ORCA-FLASH4.0 camera. Fluorescence values from individual cells were quantified using ImageJ. The reporter signal is calculated as the mean fold-change of the fluorescence ratio of YFP and mKate2 obtained for flow conditions compared to no-flow conditions, averaged across at least 30 cells from three independent experiments.

#### In situ oxygen concentration and uptake rate measurements

Oxygen concentration fields within the microfluidic channel were measured using an oxygen-sensitive fluorescent dye, tris(2,2’-bipyridyl)dichloro-ruthenium(II) hexahydrate (RTDP). RTDP was dissolved in the respective media at a final concentration of 5 mg/ml. Imaging was performed on Nikon Eclipse Ti microscope connected to a Hamamatsu ORCA-FLASH4.0 camera using a 10x objective and using a blue wavelength excitation and a far-red wavelength emission filter set. The calibration between oxygen concentration and RTDP fluorescence quenching was obtained by titrating varying amounts of dissolved oxygen in media and using the Stern-Volmer relation: *I*_0_/*I* = 1 + *K*_SV_[*O*_2_], where *I*_0_ and *I* are the fluorescence intensities without and with oxygen respectively, *K*_SV_ is the Stern-Volmer quenching constant, and [*O*_2_] is the concentration of the oxygen. All measurements including the calibration were performed within the microfluidic channel using the same illumination intensity and exposure time settings to maintain consistent fluorescence readouts. Oxygen uptake rates for each condition were calculated by measuring the average oxygen consumption over 1 h of the experiment normalized by the total number of bacterial cells.

#### Growth rate measurements

Growth rate measurements were performed using a peristaltic pump (Masterflex L/S, USA) connected to the microfluidic channel and flow within a closed-loop. Overnight-grown cells were back diluted 1:200 in fresh media and grown to an OD of 0.1 at room temperature in culture tubes with shaking. Then, the cells were loaded from the culture tubes into the channel using the pump to completely fill the channel and the connecting tubing. Thereafter, the flow loop was closed, and cells were subject to varying flow conditions for 2.5 h. After exposure to flow, the cell suspension from within the channel was collected and the final optical density (OD) at 600 nm was measured using a spectrophotometer. Growth rate was calculated according to the formula:

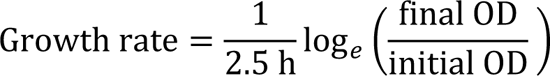

#### Reaction-diffusion transport model

To model the dynamics of oxygen concentration during flow of planktonic bacteria, we invoked the unsteady, one-dimensional reaction-diffusion model. Using the channel width coordinate *y* centered at channel middle, cell density distribution ρ(*y*), diffusivity of oxygen *D*, cellular oxygen consumption rate λ, the evolution of oxygen concentration *n*(*y, t*) in time *t* was modeled as:

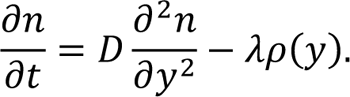

Refer to the **Supplementary Text** for details on model assumptions, derivations, and parameters. A custom MATLAB code was used to solve the above equation and simulate the evolution of oxygen and other species’ concentration fields in flow for a given experimentally-obtained distribution of planktonic bacteria.

## Supplementary Text

### Model for chemical species transport during shear flow of planktonic bacteria

We here present a continuum model based on reaction-diffusion transport to predict concentration profiles of chemical species around planktonic bacteria in shear flow. Our primary focus here is to model oxygen concentration profiles across the channel width, although a similar formulation can be used to model chemical interactions of bacteria with other chemical species in flow. Specifically, we here model the species concentration distribution across the channel width coordinate *y*, where the time *t* represents the duration under flow. Moreover, for simplicity, we assume that the cell density distribution ρ(*y*) across the channel width, which arises from motility-dependent redistribution of cells under shear flow, does not change with time. This is because cells initially redistribute over a short time scale compared to the time scales of interest, and cells do not divide and change in number significantly over the time scales of interest. Further, denoting the species diffusivity by *D*, the evolution of species concentration *n*(*y, t*) across the channel width is governed by cellular uptake and species diffusion, and is thus modeled as

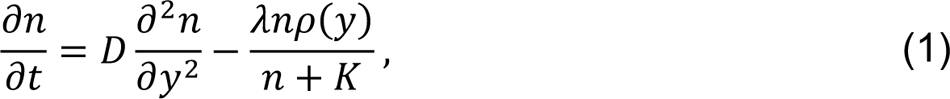

In Eq. (1), we have assumed Monod-type kinetics for species consumption by the cells, with Monod constant *K* and maximum consumption rate λ (with units mol/cell/s). In our experiments, the value of oxygen concentration in the media is in the ∼100 μM concentration range and is much higher than the Monod constant for oxygen, which is typically around ∼100 nM,(*37*) i.e., *n* ≫ *K*. Therefore, we can simplify the model in Eq. (1) as

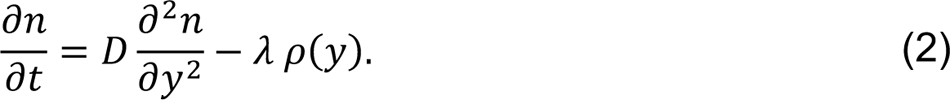

To obtain oxygen concentration profiles in Figure 3e and **Supplementary Figure 8a**, we numerically integrate Eq. (2) assuming experimentally derived values for uptake rate λ = 0.14 amol/cell/s and measured cell density distribution ρ(*y*). We assume no-flux boundary conditions at the channel walls since diffusion of oxygen across the thick PDMS channel walls precludes efficient oxygen transport over our time scales of interest. We further assume an initial condition *n*_0_ for oxygen concentration equal to 250 μM and assume an average cell density concentration ρ_0_ of 2×10^8^ cells/ml based on our experimental conditions.

Further, to predict species concentration profiles in a more general way (**Supplementary Figure 7**), we non-dimensionalize Eq. (2) by normalizing species concentration *n* by its initial value *n*_0_, cell density ρ by the average value ρ_0_, spatial coordinate *y* by channel half-width *w*, and time *t* by the diffusive time scale *w*^2^/*D*, to obtain

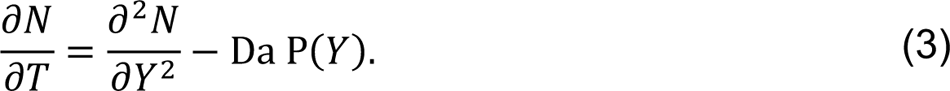

Here, 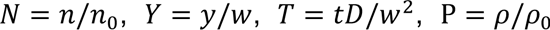, and 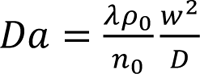 is the Damköhler number of the second kind. Da is a non-dimensional number that represents the uptake rate versus the rate of diffusion, or equivalently, the ratio of the diffusion timescale across the channel width versus the timescale of cellular uptake. Therefore, for a given uptake/consumption rate, a lower species diffusivity corresponds to a higher Da. Lastly, to obtain steady-state species concentration profiles (which is typically obtained after several diffusion time scales) in **Supplementary Figure 7**, we set the left-hand side in Eq. (3) to zero.

---

**Supplementary figure 1.**
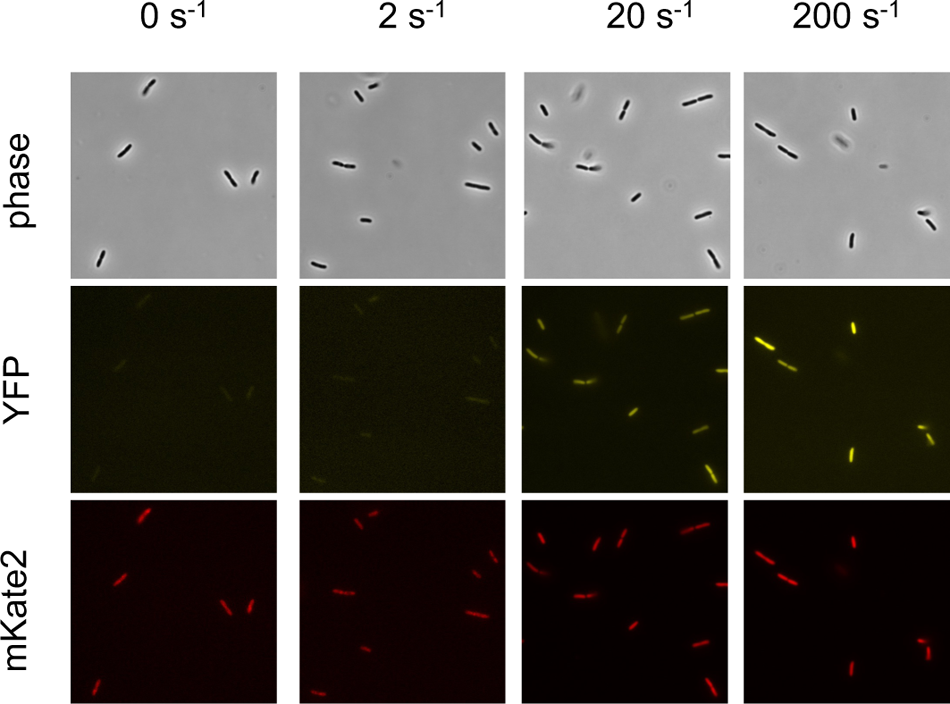
Phase contrast and fluorescence microscopy images of cells after subjecting them to flow at varying shear rates of 0, 2, 20, and 200 s^-1^ for 60 min. YFP and mKate2 represent transcriptional reporter signals from individual cells corresponding to the transcription of the *arc* operon and constitutively expressed *rpoD*, respectively.

**Supplementary figure 2.**
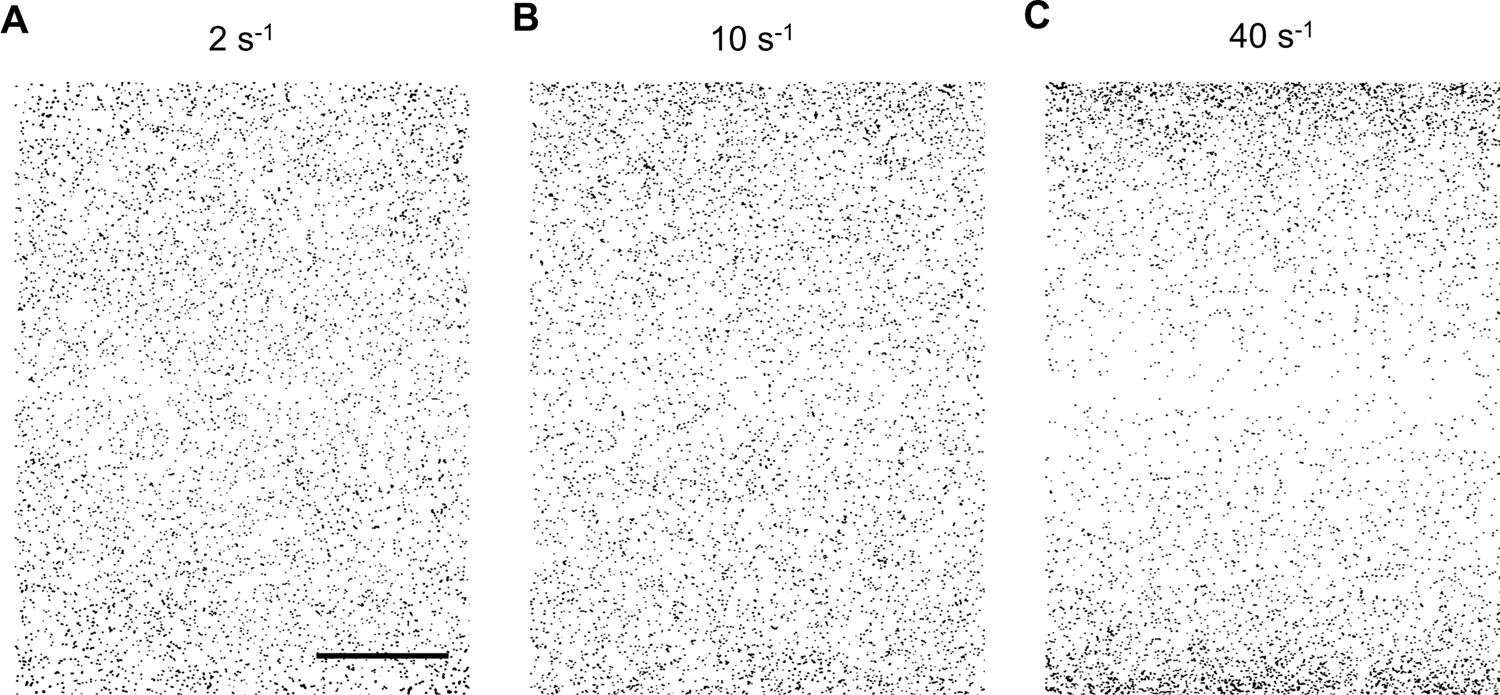
Cell distribution of WT *P. aeruginosa* measured across the channel width for shear rates of (**A**) 2 s^-1^, (**B**) 10 s^-1^, and (**C**) 40 s^-1^. Scale bar: 300 μm.

**Supplementary figure 3.**
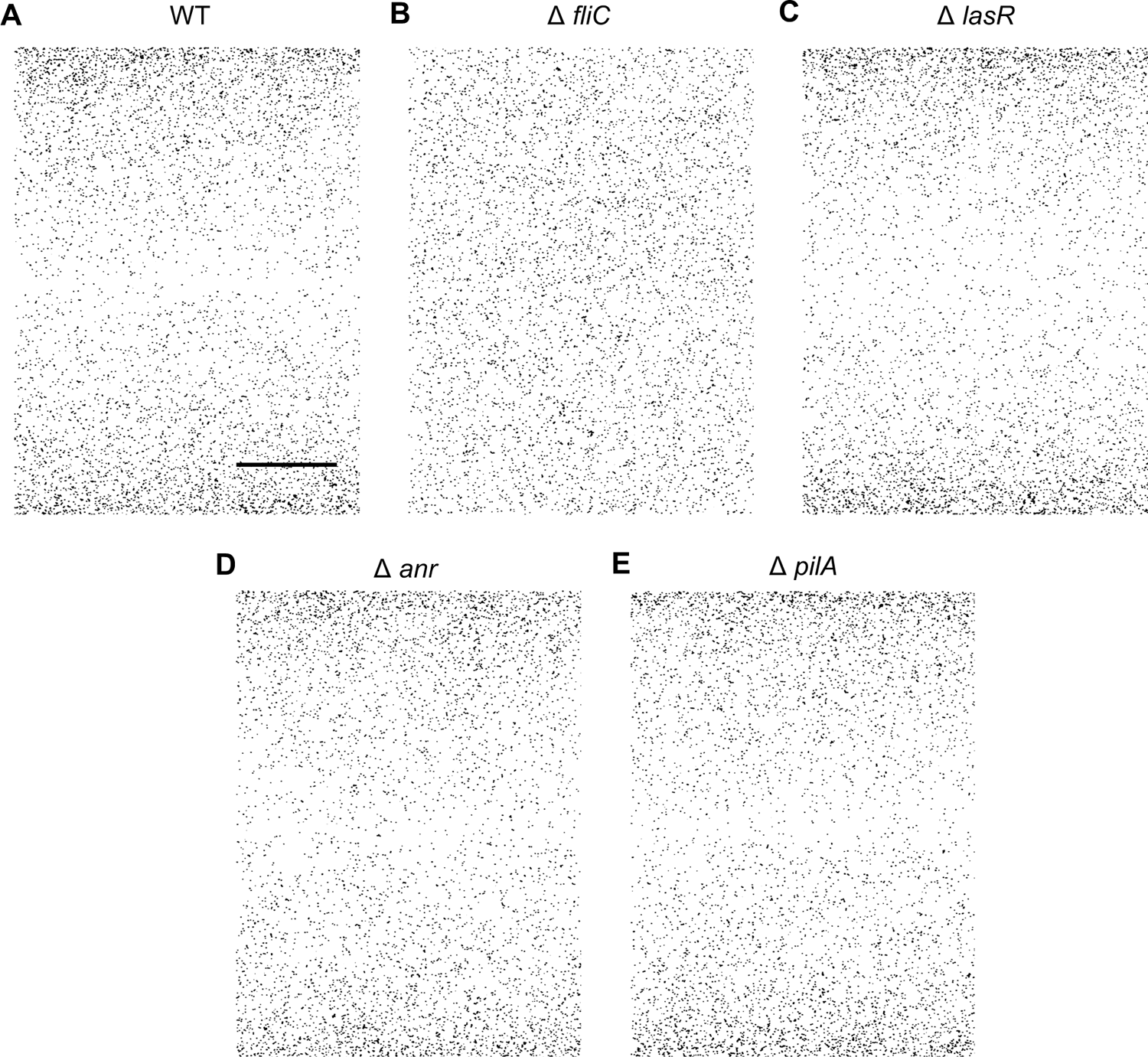
Cell distribution measured across the channel width at a shear rate of 20 s^-1^ for the following *P. aeruginosa* strains in EZ media: (**A**) WT (same as Fig. 2A), (**B**) Δ*fliC*, (**C**) Δ*lasR*, (**D**) Δ*anr*, and (**E**) Δ*pilA*. Scale bar: 300 μm.

**Supplementary figure 4.**
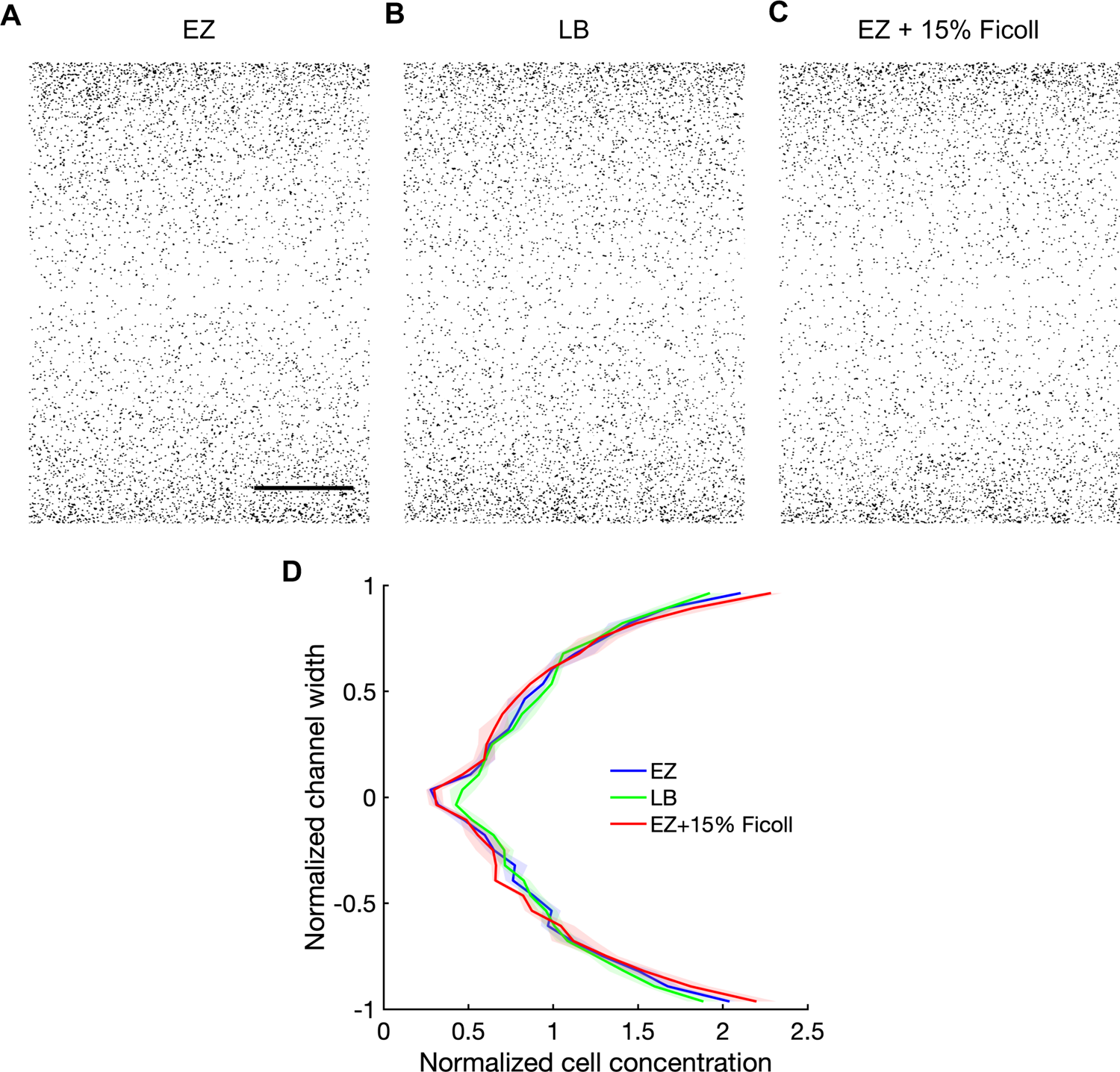
Cell distribution of WT *P. aeruginosa* measured across the channel width at a shear rate of 20 s^-1^ using (**A**) EZ rich media (same as Fig. 2A), (**B**) LB media, and (**C**) EZ rich media supplemented with 15% Ficoll. (**D**) Cell density distribution across the channel width averaged within the field of view for (**A**), (**B**), and (**C**). Scale bar: 300 μm.

**Supplementary figure 5.**
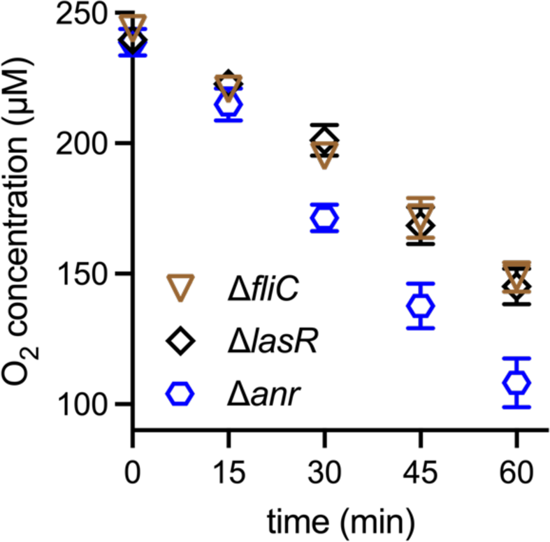
Average oxygen concentration within the field of view versus time for flow of Δ*fliC*, Δ*lasR*, and Δ*anr* strains of *P. aeruginosa* at a shear rate of 20 s^-1^. Plotted are the mean values from three independent experiments and the error bars represent the standard deviation.

**Supplementary figure 6.**
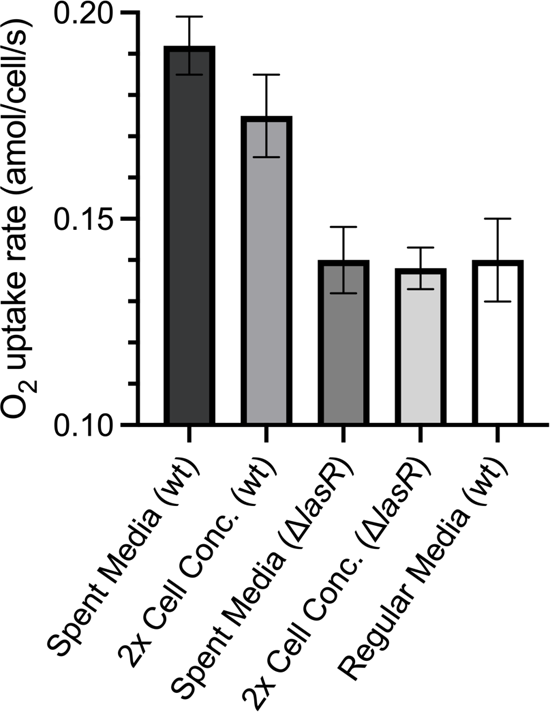
Average oxygen consumption rate of WT and Δ*lasR* strains obtained by reconstituting cells in spent media from overnight culture or by reconstituting cells at twice the initial cell concentration within the same media. Data presented here are under no-flow conditions for cells within the microfluidic channel.

**Supplementary figure 7.**
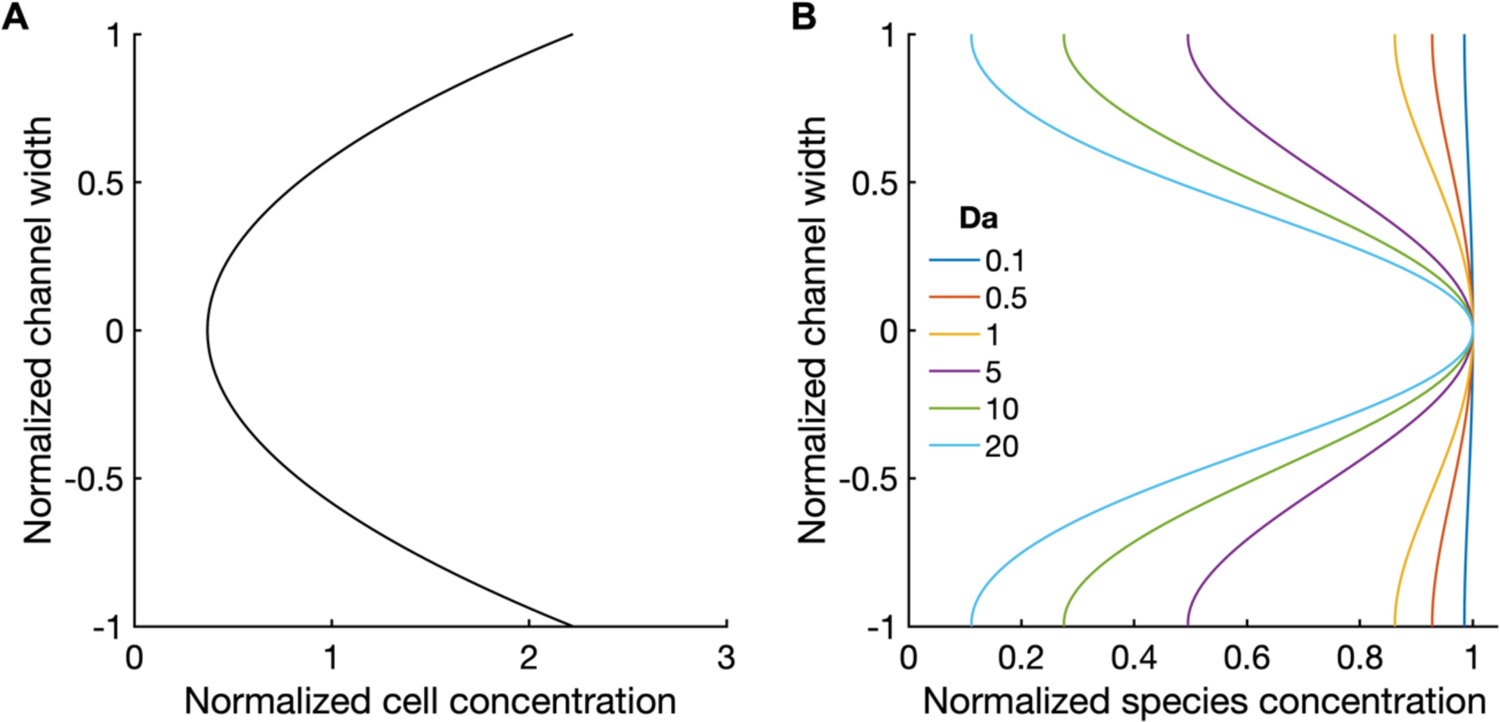
Concentration fields obtained from the reaction-diffusion model. (**A**) Approximate parabolic distribution for cell density distribution across the channel width used in the model, based on experimental observations for flow of WT *P. aeruginosa* at 20 s^-1^. (**B**) Predicted steady-state concentration fields of species for varying Damköhler number (of the second kind), Da. Da is a non-dimensional number that represents the ratio of cellular uptake/consumption rate relative to the species diffusion rate. For a given uptake rate, a decrease in species diffusivity corresponds to a proportional increase in Da.

**Supplementary figure 8.**
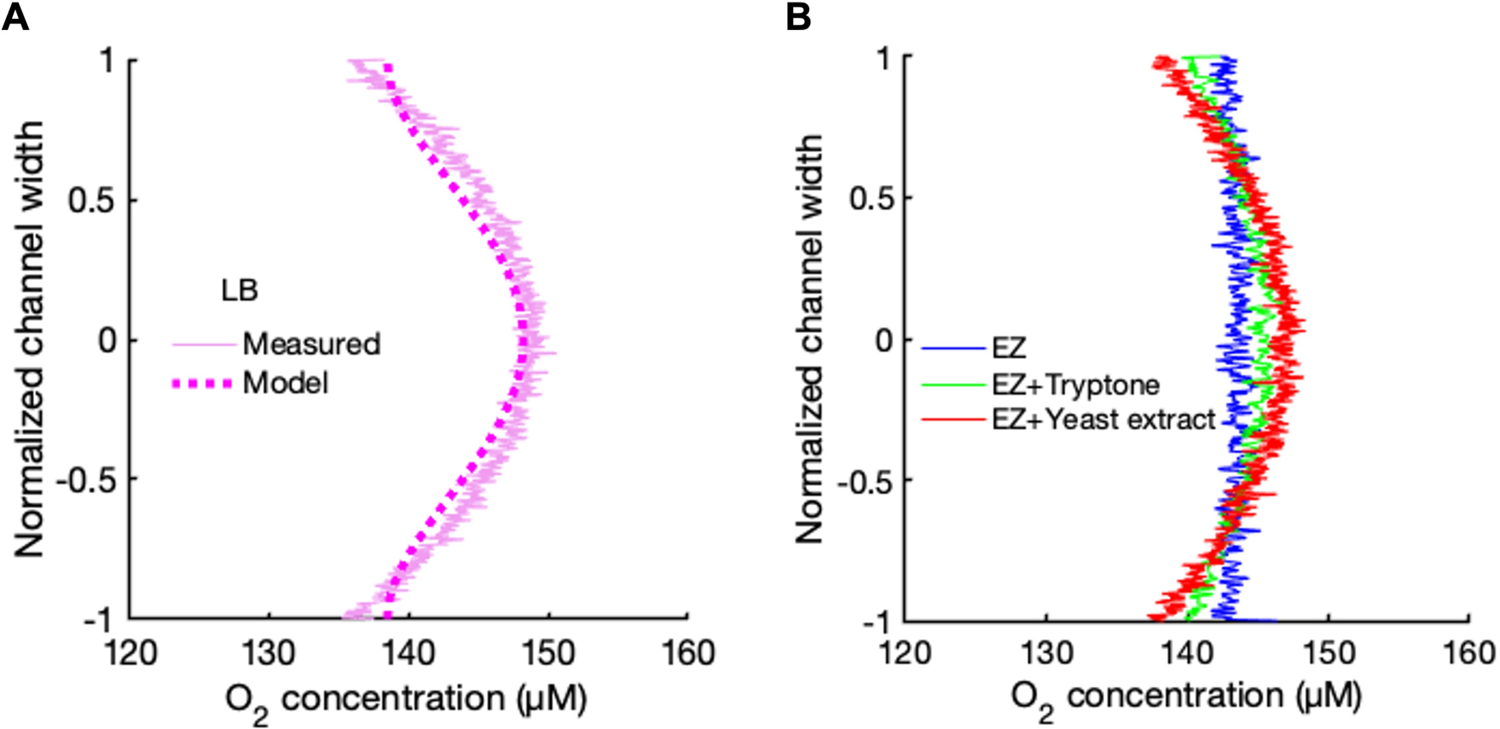
(**A**) Measured oxygen concentration (solid lines) averaged across the streamwise direction within the field of view versus channel width after 1 h of flow at shear rate of 20 s^-1^ of Δ*lasR* strain in LB media. Dashed lines represent the best fit using the reaction-diffusion model with an effective oxygen diffusivity of 0.7×10^-9^ m^2^/s. (**B**) Measured oxygen concentration profiles for similar conditions as (**A**) but using EZ media supplemented with tryptone and yeast extract in concentrations typically used for LB.

**Supplementary table 1.** Genes in planktonic PAO1 significantly upregulated by at least 3-fold in 20 s^-1^ flow compared to no flow based on RNA-Seq data. Highlighted genes correspond to marked symbols in Figure 1b which indicate genes that are typically associated with response to low oxygen conditions.

| Gene | Description | log<sub>2</sub>(fold change) |
| --- | --- | --- |
| **narI** | respiratory nitrate reductase gamma chain | 4.48 |
| **PA1746** | hypothetical protein | 4.43 |
| **narJ** | respiratory nitrate reductase delta chain | 4.31 |
| PA1137 | probable oxidoreductase | 4.27 |
| PA3871 | probable peptidyl-prolyl cis-trans isomerase, PpiC-type | 3.57 |
| **arcB** | ornithine carbamoyltransferase, catabolic | 3.28 |
| **arcC** | carbamate kinase | 3.08 |
| **narH** | respiratory nitrate reductase beta chain | 3.03 |
| PA1668 | hypothetical protein | 2.68 |
| PA0141 | conserved hypothetical protein | 2.68 |
| **arcA** | arginine deiminase | 2.55 |
| **ccoO2** | Cytochrome c oxidase, cbb3-type, CcoO subunit | 2.52 |
| **narG** | respiratory nitrate reductase alpha chain | 2.52 |
| PA1136 | probable transcriptional regulator | 2.49 |
| **ccoP2** | Cytochrome c oxidase, cbb3-type, CcoP subunit | 2.47 |
| PA1658 | conserved hypothetical protein | 2.45 |
| PA3417 | probable pyruvate dehydrogenase E1 component, alpha subunit | 2.42 |
| PA1666 | hypothetical protein | 2.39 |
| PA0087 | hypothetical protein | 2.38 |
| PA1659 | hypothetical protein | 2.38 |
| PA1665 | hypothetical protein | 2.37 |
| PA1657 | conserved hypothetical protein | 2.28 |
| PA1669 | hypothetical protein | 2.25 |
| **ccoN2** | Cytochrome c oxidase, cbb3-type, CcoN subunit | 2.24 |
| PA1662 | probable ClpA/B-type protease | 2.17 |
| PA1664 | hypothetical protein | 2.16 |
| PA1663 | probable transcriptional regulator | 2.15 |
| PA0084 | conserved hypothetical protein | 2.08 |
| PA1660 | hypothetical protein | 2.07 |
| PA1667 | hypothetical protein | 2.07 |
| **hcnB** | hydrogen cyanide synthase HcnB | 2.02 |
| PA0088 | hypothetical protein | 2.00 |
| **moaA1** | molybdopterin biosynthetic protein A1 | 1.99 |
| PA0086 | hypothetical protein | 1.90 |
| **hcnC** | hydrogen cyanide synthase HcnC | 1.88 |
| PA0089 | hypothetical protein | 1.86 |
| PA3661 | hypothetical protein | 1.84 |
| PA0083 | conserved hypothetical protein | 1.83 |
| PA3519 | hypothetical protein | 1.81 |
| PA1221 | hypothetical protein | 1.78 |
| clpV1 | ClpV1 | 1.76 |
| **hcnA** | hydrogen cyanide synthase HcnA | 1.75 |
| PA1845 | hypothetical protein | 1.74 |
| PA1844 | hypothetical protein | 1.71 |
| PA3484 | hypothetical protein | 1.67 |
| PA0098 | hypothetical protein | 1.64 |
| PA0276 | hypothetical protein | 1.62 |
| vgrG1 | VgrG1 | 1.62 |

**Supplementary table 2.** List of strains used in this study.

| Strain | Description | Reference |
| --- | --- | --- |
| ***E. coli*** |  |  |
| ZG54 | Wild-type S17 used for cloning and conjugation | Lab stock |
| ZGXXXX | S17 donor strain for $\Delta anr$ knockout in PAO1 | This study |
| ZGXXXX | S17 donor strain for $\Delta lasR$ knockout in PAO1 | This study |
| ZGXXXX | S17 maintenance strain for pArc | This study |
| ***P. aeruginosa*** |  |  |
| ZG500 | Wild-type PAO1 from Manoil transposon mutant library | Lab stock |
| ZGXXXX | PAO1 $\Delta anr$, in frame deletion of *anr* | This study |
| ZGXXXX | PAO1 $\Delta lasR$, in frame deletion of *lasR* | This study |
| ZG1593 | PAO1 $\Delta fliC$, in frame deletion of *fliC* | Lab stock |
| ZG525 | PAO1 $\Delta pilA$, in frame deletion of *pilA* | Lab stock |
| ZGXXXX | PAO1 $\Delta anr$, with pArc plasmid | This study |
| ZGXXXX | PAO1 $\Delta lasR$, with pArc plasmid | This study |
| ZGXXXX | PAO1 $\Delta fliC$, with pArc plasmid | This study |
| ZGXXXX | PAO1 $\Delta pilA$, with pArc plasmid | This study |

**Supplementary table 3.** List of plasmids used in this study.

| Plasmid | Description | Reference |
| --- | --- | --- |
| pEXG2 | Vector for generating deletion mutants | Ref. 27 |
| pEXG2-DlasR | $\Delta lasR$ knockout construct | This study |
| pEXG2-Danr | $\Delta anr$ knockout construct | This study |
| ZG1190-pPaQa | pUCP18 backbone with PPaQa::YFP, PrpoD::mKate2 | Ref. 28 |
| pArc | pUCP18 backbone with Parc::YFP, PrpoD::mKate2 | This study |

**Supplementary table 4.**
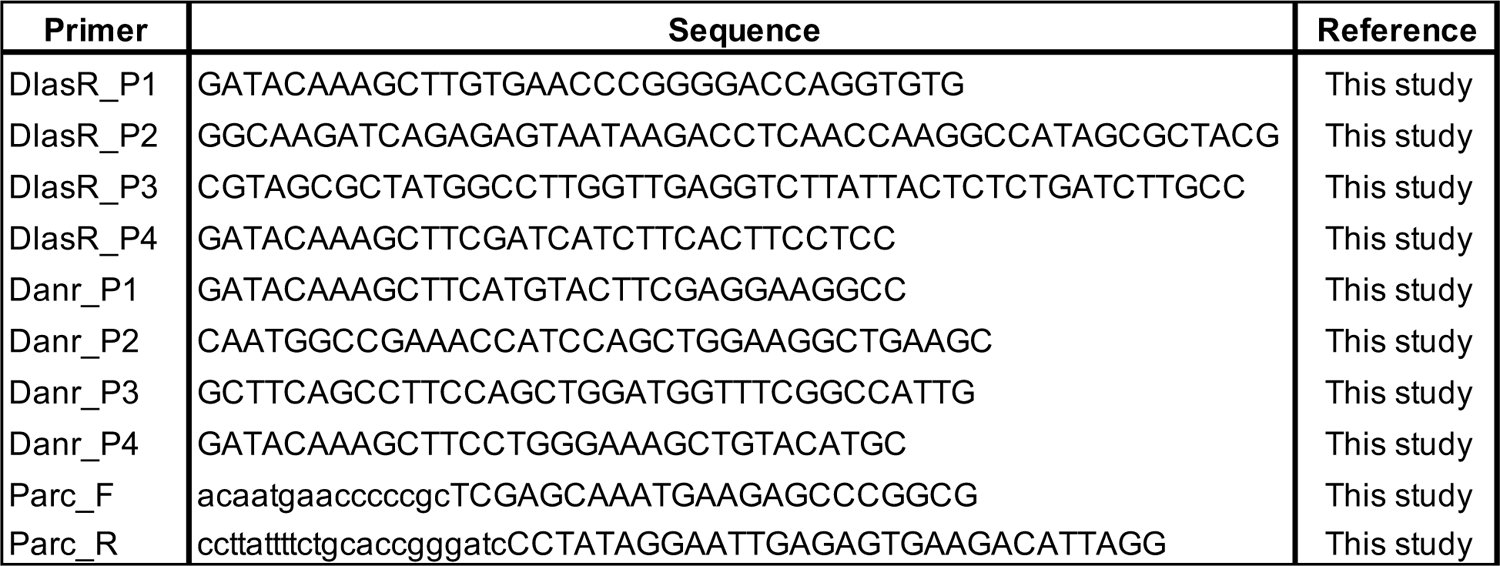
List of primers used in this study.

## References and notes

1. A. Persat, C. D. Nadell, M. K. Kim, F. Ingremeau, A. Siryaporn, K. Drescher, N. S. Wingreen, B. L. Bassler, Z. Gitai, H. A. Stone, The mechanical world of bacteria. Cell 161, 988–997 (2015).

2. Y. F. Dufrêne, A. Persat, Mechanomicrobiology: how bacteria sense and respond to forces. Nat. Rev. Microbiol. 18, 227–240 (2020).

3. B. W. Trautner, R. O. Darouiche, Catheter-Associated Infections: Pathogenesis Affects Prevention. Arch. Intern. Med. 164, 842–850 (2004).

4. H. Minasyan, Sepsis: Mechanisms of bacterial injury to the patient. Scand. J. Trauma. Resusc. Emerg. Med. 27, 1–22 (2019).

5. G. C. Padron, A. M. Shuppara, J.-J. S. Palalay, A. Sharma, J. E. Sanfilippo, Bacteria in Fluid Flow. J. Bacteriol. 205 (2023).

6. C. A. Rodesney, B. Roman, N. Dhamani, B. J. Cooley, P. Katira, A. Touhami, V. D. Gordon, Mechanosensing of shear by Pseudomonas aeruginosa leads to increased levels of the cyclic-di-GMP signal initiating biofilm development. Proc. Natl. Acad. Sci. U. S. A. 114, 5906–5911 (2017).

7. J. E. Sanfilippo, A. Lorestani, M. D. Koch, B. P. Bratton, A. Siryaporn, H. A. Stone, Z. Gitai, Microfluidic-based transcriptomics reveal force-independent bacterial rheosensing. Nat. Microbiol. 4, 1274–1281 (2019).

8. G. C. Padron, A. M. Shuppara, A. Sharma, M. D. Koch, J.-J. S. Palalay, J. N. Radin, T. E. Kehl-Fie, J. A. Imlay, J. E. Sanfilippo, Shear rate sensitizes bacterial pathogens to H 2 O 2 stress. Proc. Natl. Acad. Sci. 120, 2017 (2023).

9. T. Rossy, C. D. Nadell, A. Persat, Cellular advective-diffusion drives the emergence of bacterial surface colonization patterns and heterogeneity. Nat. Commun. 10, 1–9 (2019).

10. J. C. Conrad, R. Poling-Skutvik, Confined Flow: Consequences and Implications for Bacteria and Biofilms. Annu. Rev. Chem. Biomol. Eng 9, 175–200 (2018).

11. E. Secchi, A. Vitale, G. L. Miño, V. Kantsler, L. Eberl, R. Rusconi, R. Stocker, The effect of flow on swimming bacteria controls the initial colonization of curved surfaces. Nat. Commun. 11, 1–12 (2020).

12. R. Chawla, R. Gupta, T. P. Lele, P. P. Lele, A Skeptic’s Guide to Bacterial Mechanosensing. J. Mol. Biol. 432, 523–533 (2020).

13. J. D. Wheeler, E. Secchi, R. Rusconi, R. Stocker, Not Just Going with the Flow: The Effects of Fluid Flow on Bacteria and Plankton. Annu. Rev. Cell Dev. Biol. 35, 213–237 (2019).

14. M. M. C. Velraeds, B. Van De Belt-Gritter, H. C. Van Der Mei, G. Reid, H. J. Busscher, Interference in initial adhesion of uropathogenic bacteria and yeasts to silicone rubber by a Lactobacillus acidophilus biosurfactant. J. Med. Microbiol. 47, 1081–1085 (1998).

15. R. H. Haynes, The Rheology of Blood. Trans. Soc. Rheol. 5, 85–101 (1961).

16. C. Alvarez-Ortega, C. S. Harwood, Responses of Pseudomonas aeruginosa to low oxygen indicate that growth in the cystic fibrosis lung is by aerobic respiration. Mol. Microbiol. 65, 153–165 (2007).

17. K. Schreiber, R. Krieger, B. Benkert, M. Eschbach, H. Arai, M. Schobert, D. Jahn, The anaerobic regulatory network required for Pseudomonas aeruginosa nitrate respiration. J. Bacteriol. 189, 4310–4314 (2007).

18. G. Pessi, D. Haas, Transcriptional control of the hydrogen cyanide biosynthetic genes hcnABC by the anaerobic regulator ANR and the quorum-sensing regulators LasR and RhlR in Pseudomonas aeruginosa. J. Bacteriol. 182, 6940– 6949 (2000).

19. A. Persat, Y. F. Inclan, J. N. Engel, H. A. Stone, Z. Gitai, Type IV pili mechanochemically regulate virulence factors in Pseudomonas aeruginosa. Proc. Natl. Acad. Sci. 112, 7563–7568 (2015).

20. E. Lauga, Bacterial Hydrodynamics. Annu. Rev. Fluid Mech. 48, 105–130 (2016).

21. R. Rusconi, J. S. Guasto, R. Stocker, Bacterial transport suppressed by fluid shear. Nat. Phys. 10, 212–217 (2014).

22. L. Vennamneni, S. Nambiar, G. Subramanian, Shear-induced migration of microswimmers in pressure-driven channel flow. J. Fluid Mech. 890 (2020).

23. B. Ezhilan, D. Saintillan, Transport of a dilute active suspension in pressure-driven channel flow. J. Fluid Mech. 777, 482–522 (2015).

24. M. Gamper, A. Zimmermann, D. Haas, Anaerobic regulation of transcription initiation in the arcDABC operon of Pseudomonas aeruginosa. J. Bacteriol. 173, 4742–4750 (1991).

25. E. C. Pesci, J. P. Pearson, P. C. Seed, B. H. Iglewski, Regulation of las and rhl quorum sensing in Pseudomonas aeruginosa. J. Bacteriol. 179, 3127–3132 (1997).

26. J. Lee, L. Zhang, The hierarchy quorum sensing network in Pseudomonas aeruginosa. Protein Cell 6, 26–41 (2015).

27. L. L. Burrows, Pseudomonas aeruginosa Twitching Motility: Type IV Pili in Action. Annu. Rev. Microbiol. 66, 493–520 (2012).

28. M. Bouteiller, C. Dupont, Y. Bourigault, X. Latour, C. Barbey, Y. Konto-Ghiorghi, A. Merieau, Pseudomonas Flagella: Generalities and Specificities. Int. J. Mol. Sci. 22, 3337 (2021).

29. H. Arai, Regulation and Function of Versatile Aerobic and Anaerobic Respiratory Metabolism in Pseudomonas aeruginosa. Front. Microbiol. 2, 1–13 (2011).

30. P. W. Davenport, J. L. Griffin, M. Welch, Quorum sensing is accompanied by global metabolic changes in the opportunistic human pathogen Pseudomonas aeruginosa. J. Bacteriol. 197, 2072–2082 (2015).

31. J. H. Hammond, E. F. Dolben, T. J. Smith, S. Bhuju, D. A. Hogan, Links between Anr and quorum sensing in Pseudomonas aeruginosa biofilms. J. Bacteriol. 197, 2810–2820 (2015).

32. V. E. Wagner, D. Bushnell, L. Passador, A. I. Brooks, B. H. Iglewski, Microarray Analysis of Pseudomonas aeruginosa Quorum-Sensing Regulons: Effects of Growth Phase and Environment. J. Bacteriol. 185, 2080–2095 (2003).

33. S. C. Bryant, R. M. Navari, Effect of plasma proteins on oxygen diffusion in the pulmonary capillaries. Microvasc. Res. 7, 120–130 (1974).

34. G. S. Crowther, C. H. Chilton, S. L. Todhunter, S. Nicholson, J. Freeman, S. D. Baines, M. H. Wilcox, Comparison of planktonic and biofilm-associated communities of Clostridium difficile and indigenous gut microbiota in a triple-stage chemostat gut model. J. Antimicrob. Chemother. 69, 2137–2147 (2014).

35. F. C. Neidhardt, P. L. Bloch, D. F. Smith, Culture Medium for Enterobacteria. J. Bacteriol. 119, 736–747 (1974).

36. L. R. Hmelo, B. R. Borlee, H. Almblad, M. E. Love, T. E. Randall, B. S. Tseng, C. Lin, Y. Irie, K. M. Storek, J. J. Yang, R. J. Siehnel, P. L. Howell, P. K. Singh, T. Tolker-Nielsen, M. R. Parsek, H. P. Schweizer, J. J. Harrison, Precision-engineering the Pseudomonas aeruginosa genome with two-step allelic exchange. Nat. Protoc. 10, 1820–1841 (2015).

37. D. A. Stolpera, N. P. Revsbech, D. E. Canfield, Aerobic growth at nanomolar oxygen concentrations. Proc. Natl. Acad. Sci. U. S. A. 107, 18755–18760 (2010).

